# Plasticity in intrinsic excitability of hypothalamic magnocellular neurosecretory neurons in late-pregnant and lactating rats

**DOI:** 10.1101/2021.06.01.446490

**Authors:** Michael R. Perkinson, Rachael A. Augustine, Gregory T. Bouwer, Emily F. Brown, Isaiah Cheong, Alexander J. Seymour, Martin Fronius, Colin H. Brown

## Abstract

Oxytocin and vasopressin secretion from the posterior pituitary gland are required for normal pregnancy and lactation. Oxytocin secretion is relatively low and constant under basal conditions but becomes pulsatile during birth and lactation to stimulate episodic contraction of the uterus for delivery of the fetus and milk ejection during suckling. Vasopressin secretion is maintained in pregnancy and lactation despite reduced osmolality (the principal stimulus for vasopressin secretion) to increase water retention to cope with the cardiovascular demands of pregnancy and lactation. Oxytocin and vasopressin secretion are determined by the action potential (spike) firing of magnocellular neurosecretory neurons of the hypothalamic supraoptic and paraventricular nucleus. In addition to synaptic input activity, spike firing depends on intrinsic excitability conferred by the suite of channels expressed by the neurons. Therefore, we analysed oxytocin and vasopressin neuron activity in anaesthetised non-pregnant, late-pregnant and lactating rats to test the hypothesis that intrinsic excitability of oxytocin and vasopressin neurons is increased in late pregnancy and lactation to promote oxytocin and vasopressin secretion required for successful pregnancy and lactation. Hazard analysis of spike firing revealed a higher incidence of post-spike hyperexcitability immediately following each spike in oxytocin neurons, but not in vasopressin neurons, in late pregnancy and lactation, which is expected to facilitate high frequency firing during bursts. Despite lower osmolality in late-pregnant and lactating rats, vasopressin neuron activity was not different between non-pregnant, late-pregnant and lactating rats, and blockade of osmosensitive ΔN-TRPV1 channels inhibited vasopressin neurons to a similar extent in non-pregnant, late-pregnant and lactating rats. Furthermore, supraoptic nucleus ΔN-TRPV1 mRNA expression was not different between non-pregnant and late-pregnant rats, suggesting that enhanced activity of ΔN-TRPV1 channels might maintain vasopressin neuron activity to increase water retention during pregnancy and lactation.

## Introduction

Secretion of the posterior pituitary hormones, oxytocin and vasopressin (the antidiuretic hormone), is required for successful pregnancy and lactation. Oxytocin is required for normal delivery during birth and is essential for milk-ejection during lactation (1), while vasopressin increases water retention during pregnancy for adequate placental blood supply (2). Vasopressin secretion is principally stimulated by body fluid osmolality to promote renal water reabsorption and so lowered osmolality normally decreases vasopressin secretion (3). However, the osmotic threshold for vasopressin secretion is decreased during pregnancy to increase renal water reabsorption despite lowered osmolality, to support blood volume expansion to cope with the cardiovascular demands of the developing fetus (4, 5).

Oxytocin and vasopressin are synthesised by magnocellular neurosecretory neurons of the hypothalamic supraoptic and paraventricular nuclei that each project a single axon to the posterior pituitary gland where hormone secretion into the circulation is triggered by action potential (spike) firing (1). Oxytocin neurons typically fire spikes in a slow continuous pattern (1) to maintain relatively constant oxytocin concentrations (6). However, oxytocin neurons fire short high frequency bursts every 5 – 10 min during birth and suckling (7–9) to secrete large pulses of oxytocin that induce rhythmic uterine contraction during birth and episodic milk ejection during suckling.

While afferent inputs are necessary for osmotic activation of vasopressin neurons, direct osmotically-induced depolarisation is also necessary to increase the excitability of the neurons for spike firing (10, 11). Osmotically-induced depolarisation results from activation of N-terminal truncated TRPV1 (ΔN-TRPV1) channels (12) that are mechanically gated by osmotically-induced cell shrinkage (13, 14). In addition to afferent inputs and baseline membrane potential, the excitability of magnocellular neurons depends on post-spike potentials (afterpotentials), including the fast, medium and slow afterhyperpolarisation (fAHP, mAHP and sAHP) (15–17), and the fast and slow afterdepolarisation (fADP and sADP) (18–20) that transiently modulate excitability after each spike (post-spike excitability).

Here, we analysed basal spike patterning in oxytocin and vasopressin neurons of non-pregnant, late-pregnant and lactating rats to determine whether their post-spike excitability is increased in pregnancy and lactation. In addition, we used pharmacological blockade of ΔN-TRPV1 channels to determine whether direct osmosensitivity of vasopressin neurons is increased in late pregnancy and lactation.

## Methods

### Animals

All experimental procedures were approved by the University of Otago Animal Ethics Committee and were carried out in accordance with the New Zealand Animal Welfare Act (1999) and associated guidelines.

Animals were purchased from the University of Otago Animal Facility. Non-pregnant rats were group-housed while late-pregnant and lactating rats were individually caged from mid-pregnancy onwards, under controlled conditions (12 h light/12 h dark cycle with lights on at 07.00 h; 22 ± 1°C) with free access to food and water. To generate late-pregnant and lactating rats, the oestrous cycle was monitored daily using vaginal cytology and the rats were placed overnight in a cage with a male on pro-oestrus. The presence of sperm in the vaginal lavage on the following morning indicated mating had occurred (day 0 of gestation: G0).

Non-pregnant rats were virgin rats that were freely-cycling to prevent any potential confounding effects of the oestrous cycle on neuronal activity. Late-pregnant rats were used on G18-21. Lactating rats gave birth on G21 or 22 (day 0 post-partum, PP0) and were used on post-partum days PP07-17.

### In vivo *electrophysiology*

The rats were anaesthetised with IP urethane (1.25 – 1.50 g kg^−1^ ethyl carbamate, Sigma) for extracellular single-unit recording of spike firing. Pups were removed from lactating rats at least 4 h before experiments commenced. Upon complete cessation of the flexor withdrawal reflex, the left femoral vein was catheterised and a blood sample removed to measure plasma osmolality by freezing point depression osmometry (Model 3320; Advanced Instruments, MA, USA). Plasma osmolality was only measured in rats that received exactly 1.25 g kg^−1^ urethane because urethane is osmotically active. The rats were placed supine in a stereotaxic frame and the pituitary stalk and the right supraoptic nucleus were exposed through the oral and nasal cavities (21).

Following the removal of the meninges overlying the supraoptic nucleus, a U-shaped microdialysis probe (22), permeable to 10 kDa (Spectra/Por RC Hollow Fibers, Spectrum Medical Inc., Houston, TX, USA) was bent to position the loop of the membrane over the ventral surface of the supraoptic nucleus. The supraoptic nucleus was continuously dialysed with artificial cerebrospinal fluid (aCSF; composition in mM: NaCl 138, KCl 3.36, NaHCO3 9.52, Na_2_HPO_4_ 0.49, urea 2.16, CaCl_2_ 1.26, MgCl_2_ 1.18) at 3 μl min^−1^.

Extracellular single-unit recordings were made via a glass recording microelectrode (15 – 40 MΩ) filled with 0.9% saline placed through the loop of the microdialysis probe, using a Neurolog system connected to a CED 1401 analogue-digital interface (Cambridge Electronic Design) and Spike 2 software (Cambridge Electronic Design). Antidromic spikes were evoked in supraoptic nucleus neurons by a side-by-side SNEX-200 stimulating electrode (Science Products GmbH) placed on the pituitary stalk. The rats were euthanised by IV anaesthetic overdose or IV 3 M KCl (0.5 ml) at the end of the experiments.

Neurons that fired less than one spontaneous spike every 10 s were categorized as silent and were not recorded. Phasic activity was defined as 95% of spikes begin portioned into bursts using the following parameters: active periods lasting ≥5 s containing ≥20 spikes and ≥5 s interval between active periods during which there was ≤1 spike every 5 s; (23). Non-phasic neurons were characterised as oxytocin neurons on the basis of a transient excitation ≥0.5 spikes s^−1^ over 5 min following IV injection of (Tyr[SO_3_H]^27^)cholecystokinin fragment 26-33 amide (CCK8S; Sigma; 20 µg kg^−1^, 0.5 ml kg^−1^ in 0.9% saline), or as vasopressin neurons by transient inhibition or no change in firing rate after CCK8S (23).

Spontaneous spike firing of each neuron was recorded for at least 10 min before CCK8S administration and the first 10 min of each recording was used for analyses of activity patterning; these recordings include the baseline recording periods from neurons that were subsequently challenged with drug administration that have been previously published for other purposes (23–27). For experiments to determine the ΔN-TRPV1 contribution to basal activity, the dialysate was switched to aCSF containing 10 mM ruthenium red (Sigma) for 60 min, at least 10 min after recovery from any effects of IV CCK8S.

### Hazard function

Inter-spike interval histograms were constructed (in 0.01 s bins) for the first 10 min of recording to generate hazards for each neuron. Hazards were calculated using the formula, hazard_[i-1, i]_ = n_[i-1, i]_ / (N – n_[0, i-1]_); where hazard_[i-1, i]_ is the probability of the next spike firing in a given interval, i, n_[i-1, i]_ is the number of spikes in interval i, N is the total number of spikes in all intervals and n_[0, i-1]_ is the total number of spikes in all intervals preceding the current interval, i. The peak early hazard was taken as the maximum hazard within the first 0.07 s after each spike and the mean late hazard was calculated from the 0.41 – 0.5 s intervals after each spike. The peak early:mean late hazard ratio (hazard ratio) was then calculated.

### Post-spike hyperexcitability

To determine whether individual oxytocin neurons displayed elevated post-spike hyperexcitability within the first 0.07 s after each spike, two standard deviations of the mean hazard ratio was calculated for oxytocin neurons in non-pregnant rats and added to the mean hazard ratio of oxytocin neurons in non-pregnant rats to set the threshold to be classified as post-spike hyperexcitability for oxytocin neurons in non-pregnant, late-pregnant and lactating rats. The same procedure was applied to determine any differences in post-spike hyperexcitability for individual vasopressin neurons in non-pregnant, late-pregnant and lactating rats, using vasopressin neurons from non-pregnant rats as baseline.

### Coefficient of variation

The coefficient of variation of inter-spike interval (CV) was calculated as a measure of variability of spike firing using the formula, CV = standard deviation/mean.

### RNAscope

ΔN-TRPV1 mRNA expression was measured in supraoptic nucleus vasopressin neurons using RNAscope. Rats were deeply anaesthetised with pentobarbital (300 mg kg^−1^, IP) and perfused with 50 ml saline followed by 200 ml 4% paraformaldehyde (PFA) in 0.1 M phosphate buffer (PB). Brains were removed, post-fixed overnight in 4% PFA, transferred to 30% sucrose solution in 0.1 M PB and stored at 4°C until they were infiltrated with sucrose. Brains were then frozen on dry ice and stored at −80°C until 15 µm coronal sections were cut and slide mounted on a cryostat.

Sections were pre-treated according to the Advance Cell Diagnostics, Inc (ACD bio) RNAscope Multiplex Fluorescent Assay protocol. Briefly, sections were dehydrated through an ethanol series at room temperature (RT) and air-dried. Endogenous peroxidase activity was blocked with hydrogen peroxide (ACD bio Cat. No.323100) for 10 min at RT. Target retrieval was performed in boiling Target Retrieval Reagent (ACD bio Cat. No. 323100) for 5 min. Sections were treated with protease III (ACD bio Cat. No. 323100) for 30 min at 40°C to increase cell permeability.

Dual fluorescent RNAscope was performed using the RNAscope Multiplex Fluorescent Reagent Kit v2 (ACD bio] Cat. Nos. 323100 and 323120) according to the manufacturers protocol. Briefly, target probes for rat ΔN-TRPV1 mRNA (C2; 472 - 1431 bp, ascension no.: NM_031982.1), ACD bio) and rat vasopressin mRNA (C4; 20 – 525 bp, ascension no.: NM_016992.2) were mixed, warmed for 10 min at 40°C, and allowed to cool to RT. Sections were subjected to probe hybridisation (2 h at 40°C) and amplifier hybridisation (AMP1, AMP2, AMP3; 30min at 40°C/AMP). After washing, C2 probe signal was developed with HRP-C2 (15 min at 40°C), Opal 570 dye (Akoya Biosciences; 1:1500; 30 min at 40°C), followed by HRP blocker (15 min at 40°C). After further washing, the C4 probe signal was developed as above but with HRP-C4 and Opal 620 dye (Akoya Biosciences; 1:1500). Slides were coverslipped with ProLong Gold Antifade Mountant (Invitrogen™) and stored at 4°C.

Sections were imaged in Z-stacks at 570 nm for ΔN-TRPV1 mRNA and 620 nm for vasopressin mRNA on a confocal microscope (Nikon A1R MP). Stacks were compressed using ImageJ Fiji Z-projection Max intensity. Max intensity images were processed using Cell Profiler™ cell image analysis software to create binary threshold images (Supplementary Figure 1). The area (in pixels) of ΔN-TRPV1 mRNA co-localised with vasopressin mRNA was divided by the total area of vasopressin mRNA to give the percentage of vasopressin mRNA co-labelled with ΔN-TRPV1 mRNA, and area of ΔN-TRPV1 mRNA co-localised with vasopressin mRNA was divided by the total area of ΔN-TRPV1 mRNA to give the percentage of ΔN-TRPV1 mRNA co-labelled with vasopressin mRNA.

### Statistics

All values are reported as mean ± standard error of the mean. Statistical analyses were completed using Sigma Plot version 14 for Windows (Systat Software, San Jose, CA, USA).

## Results

### Basal firing rate of oxytocin and vasopressin neurons in non-pregnant, late-pregnant and lactating rats

Spontaneous activity was analysed from the first 10 min of 94 recordings from oxytocin neurons and 204 recordings from vasopressin neurons in 76 non-pregnant rats, 74 late-pregnant rats and 35 lactating rats. There was no effect of reproductive status on the basal firing rate of oxytocin neurons (one-way ANOVA on ranks; H = 0.35, P = 0.84) or vasopressin neurons (H = 0.56, P = 0.76; Figure 1).

**Figure 1.**
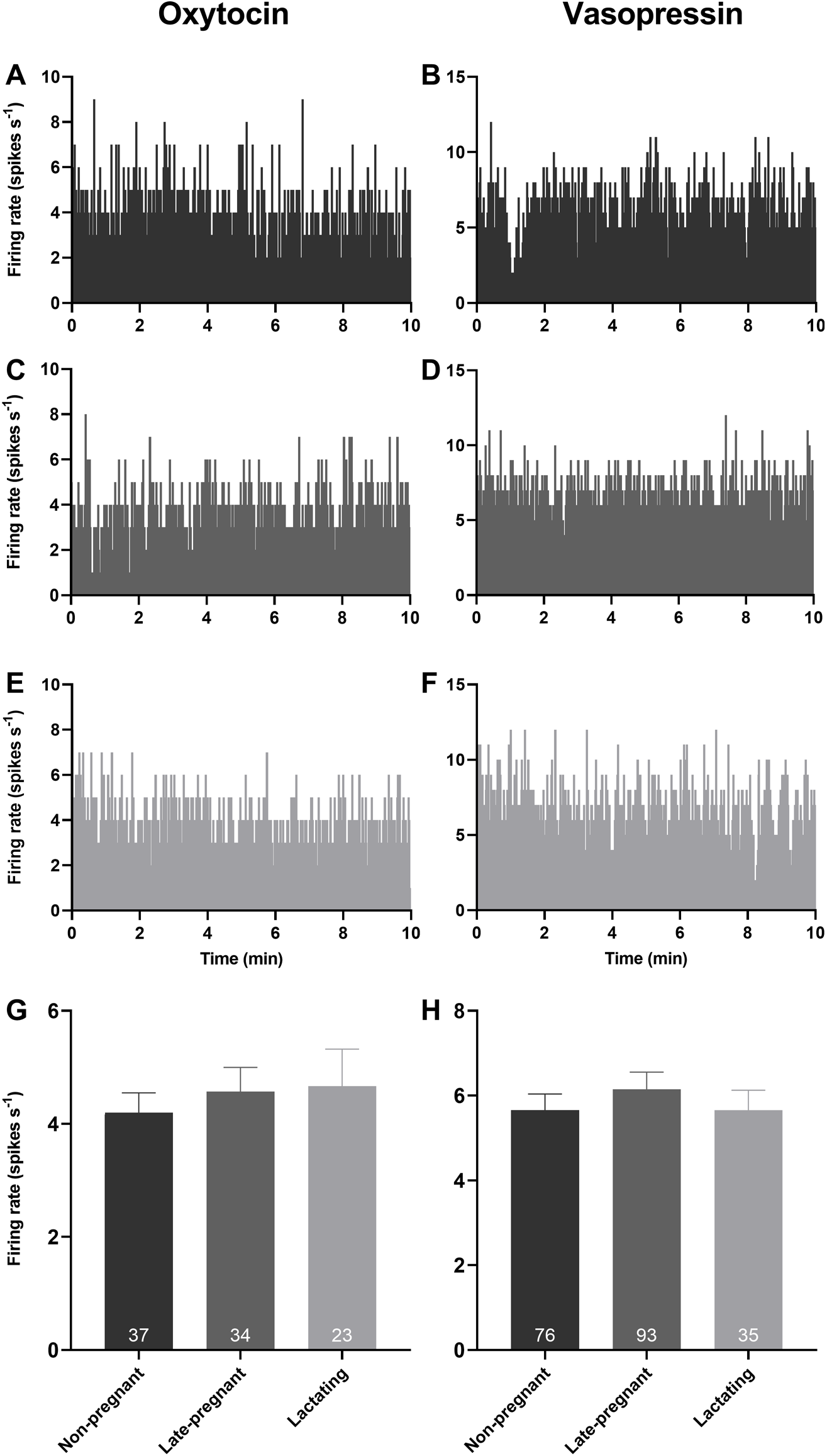
Basal firing rate of oxytocin and vasopressin neurons are not affected by reproductive status. **A – F**. Example ratemeter recordings of oxytocin neuron (A, C, E) and vasopressin neuron (B, D, F) firing rate (in 1 s bins) from urethane-anaesthetised non-pregnant (A, B), late-pregnant (C, D) and lactating (E, F) rats. **G and H**. Mean (± SEM) spontaneous oxytocin neuron (G) and vasopressin neuron (H) firing rate (averaged over 10 min) in non-pregnant, late-pregnant and lactating rats. There was no effect of reproductive status on the basal firing rate of oxytocin neurons (one-way ANOVA on ranks; H = 0.35, P = 0.84) or vasopressin neurons (H = 0.56, P = 0.76).

### Phasic activity of vasopressin neurons in non-pregnant, late-pregnant and lactating rats

Spontaneous phasic activity was evident in 54 vasopressin neurons. There was no effect of reproductive status on the overall firing rate (H = 3.52, P = 0.17), intraburst firing rate (H = 2.31, P = 0.32), burst duration (H = 0.26, P = 0.88) or interburst interval of phasic neurons (H = 4.72, P = 0.09; Figure 2).

**Figure 2.**
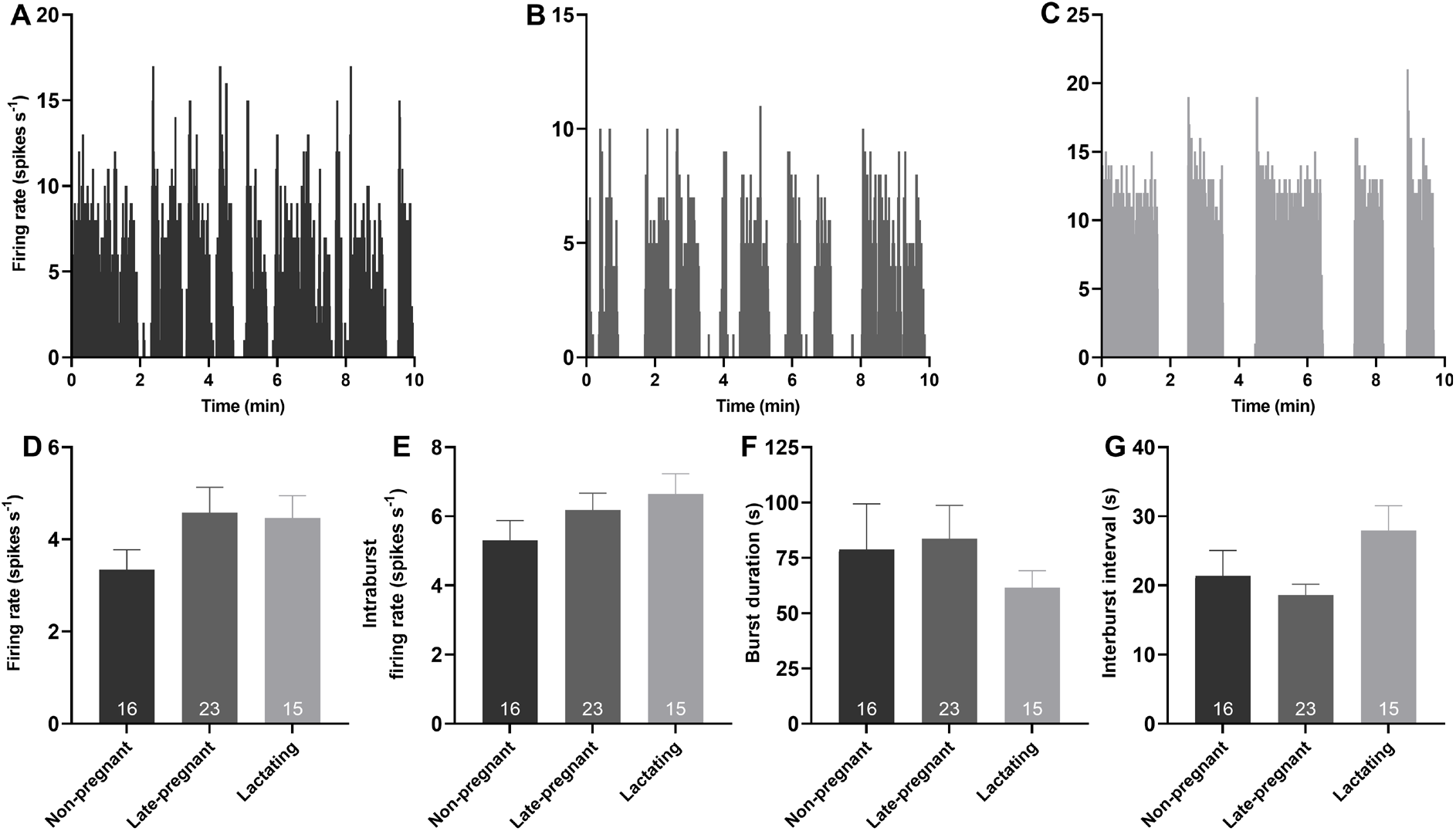
Phasic activity of vasopressin neurons is not affected by reproductive status. **A – C**. Example ratemeter recordings of spontaneous phasic activity in vasopressin neurons (in 1 s bins) from urethane-anaesthetised non-pregnant (A), late-pregnant (B) and lactating (C) rats. **D – G**. Mean (± SEM) phasic neuron firing rate (D), intraburst firing rate (E), burst duration (F) and interburst interval (G) in non-pregnant, late-pregnant and lactating rats. There was no effect of reproductive status on the firing rate (H = 3.52, P = 0.17), intraburst firing rate (H = 2.31, P = 0.32), burst duration (H = 0.26, P = 0.88) or interburst interval of phasic neurons (H = 4.72, P = 0.09).

### Post-spike excitability in oxytocin and vasopressin neurons in non-pregnant, late-pregnant and lactating rats

Hazard functions were generated to determine whether post-spike excitability of oxytocin and vasopressin neurons was different between non-pregnant, late-pregnant and lactating rats. There was no effect of reproductive status on peak early hazard (H = 1.00, P = 0.61; one-way ANOVA on ranks), mean late hazard (H = 0.09, P = 0.95) or hazard ratio (H = 1.69, P = 0.43) of oxytocin neurons (Figure 3). Nevertheless, oxytocin neurons in late-pregnant and lactating rats had a higher incidence of hazard ratios that reached the threshold to be classified as elevated post-spike hyperexcitability (χ^2^ = 7.07, P = 0.03; Figure 3F).

**Figure 3.**
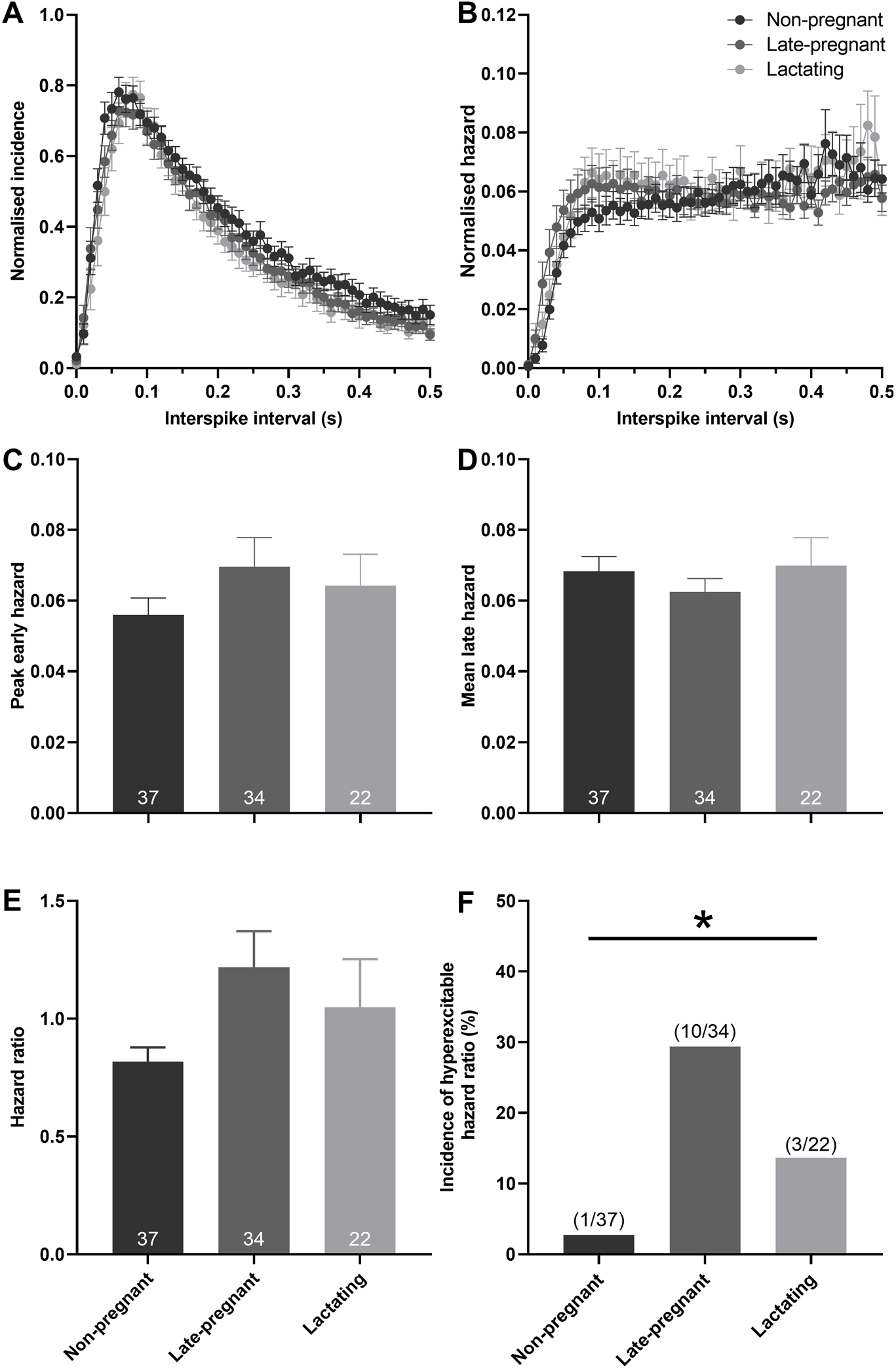
Post-spike excitability in oxytocin neurons in non-pregnant, late-pregnant and lactating rats. **A**. Mean normalised interspike intervals (± SEM) of basal activity in oxytocin neurons from non-pregnant, late-pregnant and lactating rats. **B**. Mean (± SEM) normalised hazard functions from the same neurons in A. **C - E**. Mean (± SEM) peak early hazard (C), late hazard (D) and early:mean late hazard ratio (E) in oxytocin neurons from non-pregnant, late-pregnant and lactating rats. There was no effect of reproductive status on peak early hazard (H = 1.00, P = 0.61; one-way ANOVA on ranks), mean late hazard (H = 0.09, P = 0.95) or hazard ratio (H = 1.69, P = 0.43) of oxytocin neurons. **F**. Proportion of oxytocin neurons with post-spike hyperexcitability in non-pregnant, late-pregnant and lactating rats. The incidence of post-spike hyperexcitability was different between oxytocin neurons from non-pregnant, late-pregnant and lactating rats (χ^2^ = 7.07, P = 0.03).

There was no effect of reproductive status on the peak early hazard (H = 0.82, P = 0.66), mean late hazard (H = 1.81, P = 0.40) or hazard ratio (H = 4.72, P = 0.09) of vasopressin neurons (Figure 4). Similarly, there was no effect of reproductive status on the peak early hazard (H = 0.17, P = 0.92), mean late hazard (H = 4.27, P = 0.12) or hazard ratio (H = 0.41, P = 0.82) of phasic vasopressin neurons (Supplementary Figure 2). There was no difference in the incidence of hazard ratios that reached the threshold to be classified as elevated post-spike hyperexcitability in vasopressin neurons in non-pregnant, late-pregnant and lactating rats (χ2 = 2.38, P = 0.30; Figure 4F).

**Figure 4.**
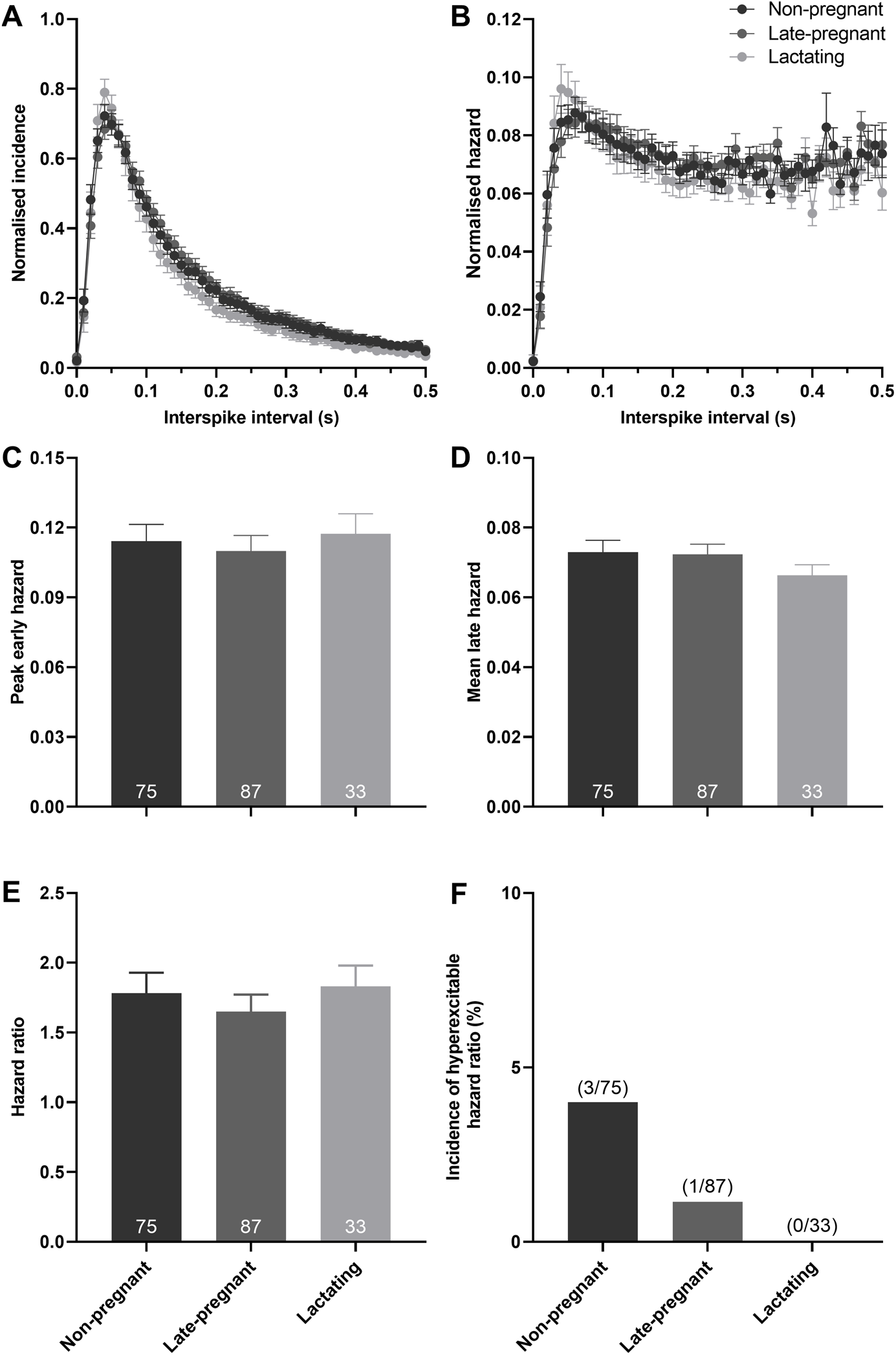
Post-spike excitability in vasopressin neurons in non-pregnant, late-pregnant and lactating rats. **A**. Mean (± SEM) normalised interspike intervals of basal activity in vasopressin neurons from non-pregnant, late-pregnant and lactating rats. **B**. Mean (± SEM) normalised hazard functions from the same neurons in A. **C - E**. Mean (± SEM) peak early hazard (C), late hazard (D) and early:mean late hazard ratio (E) in vasopressin neurons from non-pregnant, late-pregnant and lactating rats. There was no effect of reproductive status on the peak early hazard (H = 0.82, P = 0.66), mean late hazard (H = 1.81, P = 0.40) or hazard ratio (H = 4.72, P = 0.09) in vasopressin neurons. **F**. Proportion of vasopressin neurons with post-spike hyperexcitability in non-pregnant, late-pregnant and lactating rats. The incidence of post-spike hyperexcitability was not different between vasopressin neurons from non-pregnant, late-pregnant and lactating rats (χ^2^ = 2.38, P = 0.30).

### Variability of spike firing in oxytocin and vasopressin neurons in non-pregnant, late-pregnant and lactating rats

The CV was calculated to determine whether the higher incidence of post-spike hyperexcitability in oxytocin neurons of late-pregnant and lactating rats was associated with increased variability of spike firing. While there was no effect of reproductive status on the CV of oxytocin neurons (H = 0.76, P = 0.69), vasopressin neurons (H = 5.43, P = 0.07), or phasic vasopressin neurons (H = 2.73, P = 0.25; Supplementary Figure 3), there was a significant correlation between the CV and hazard ratio for oxytocin neurons (Spearman rank order correlation coefficient, r = 0.24, P = 0.03), vasopressin neurons (r = 0.49, P < 0.001) and for phasic vasopressin neurons (r = 0.43, P = 0.002; Figure 5).

**Figure 5.**
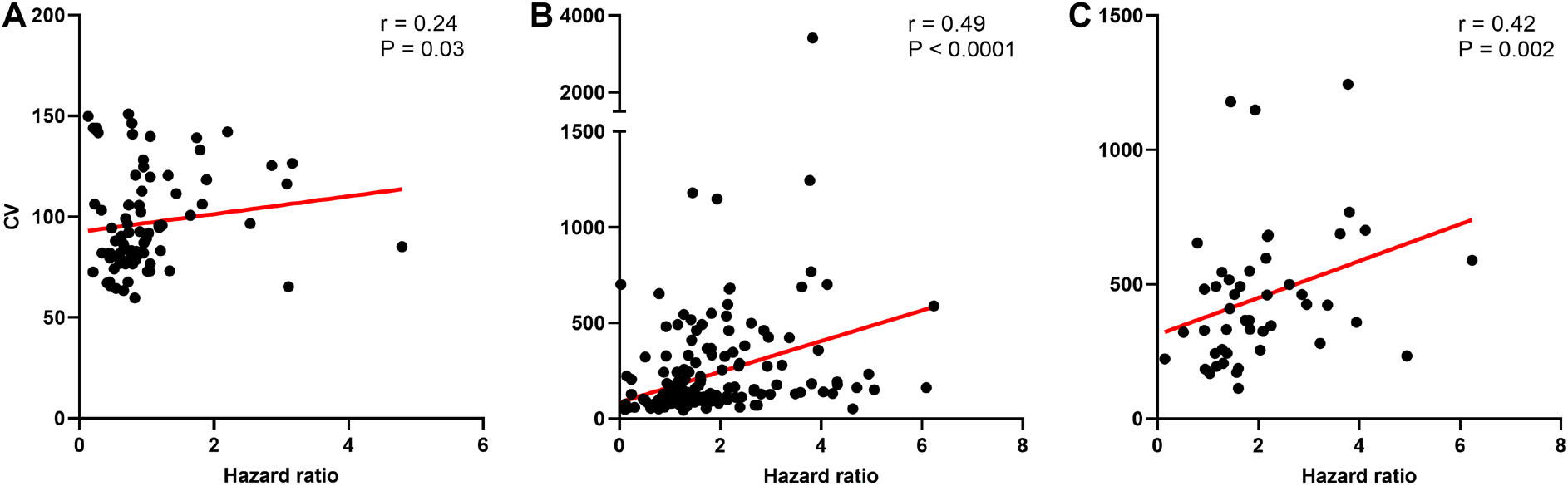
Variability of spike firing in oxytocin and vasopressin neurons in non-pregnant, late-pregnant and lactating rats. **A – C**. Scatter plots of the coefficient of variation of inter-spike interval (CV) against peak early:mean late hazard ratio in oxytocin neurons (A), vasopressin neurons (B) and phasic vasopressin neurons (C) from non-pregnant, late-pregnant and lactating rats. There was a significant correlation between the CV and hazard ratio for oxytocin neurons (Spearman rank order correlation coefficient, r = 0.24, P = 0.03), for vasopressin neurons (r = 0.49, P < 0.001) and for phasic vasopressin neurons (r = 0.43, P = 0.002). Regression lines are shown for illustrative purpose only.

### Ruthenium red effects on oxytocin and vasopressin neuron activity in non-pregnant, late-pregnant and lactating rats

As expected, plasma osmolality was affected by reproductive status (H = 20.0, P < 0.001), with lower osmolality in late-pregnant rats (P < 0.001, Dunn’s post hoc test) and lactating rats (P = 0.04) than in non-pregnant rats. Despite lower osmolality in late-pregnant and lactating rats, microdialysis administration of ruthenium red into the supraoptic nucleus reduced the firing rate of vasopressin neurons to a similar extent in non-pregnant, late-pregnant and lactating rats (F_2,28_ = 0.69, P = 0.51 for the main effect of REPRODUCTIVE STATUS; F_6,28_ = 5.94, P < 0.001 for the main effect of TIME; F_12,32_ = 0.29, P = 0.99 for the interaction between REPRODUCTIVE STATUS and TIME, two-way repeated measures ANOVA; Figure 6). There was no effect of ruthenium red on the peak early hazard (F_1,28_ = 1.70, P = 0.20), mean late hazard (F_1,26_ = 0.37, P = 0.85) or hazard ratio (F_1,26_ = 0.21, P = 0.81) of vasopressin neurons and no interaction of ruthenium red with reproductive status (Figure 6).

**Figure 6.**
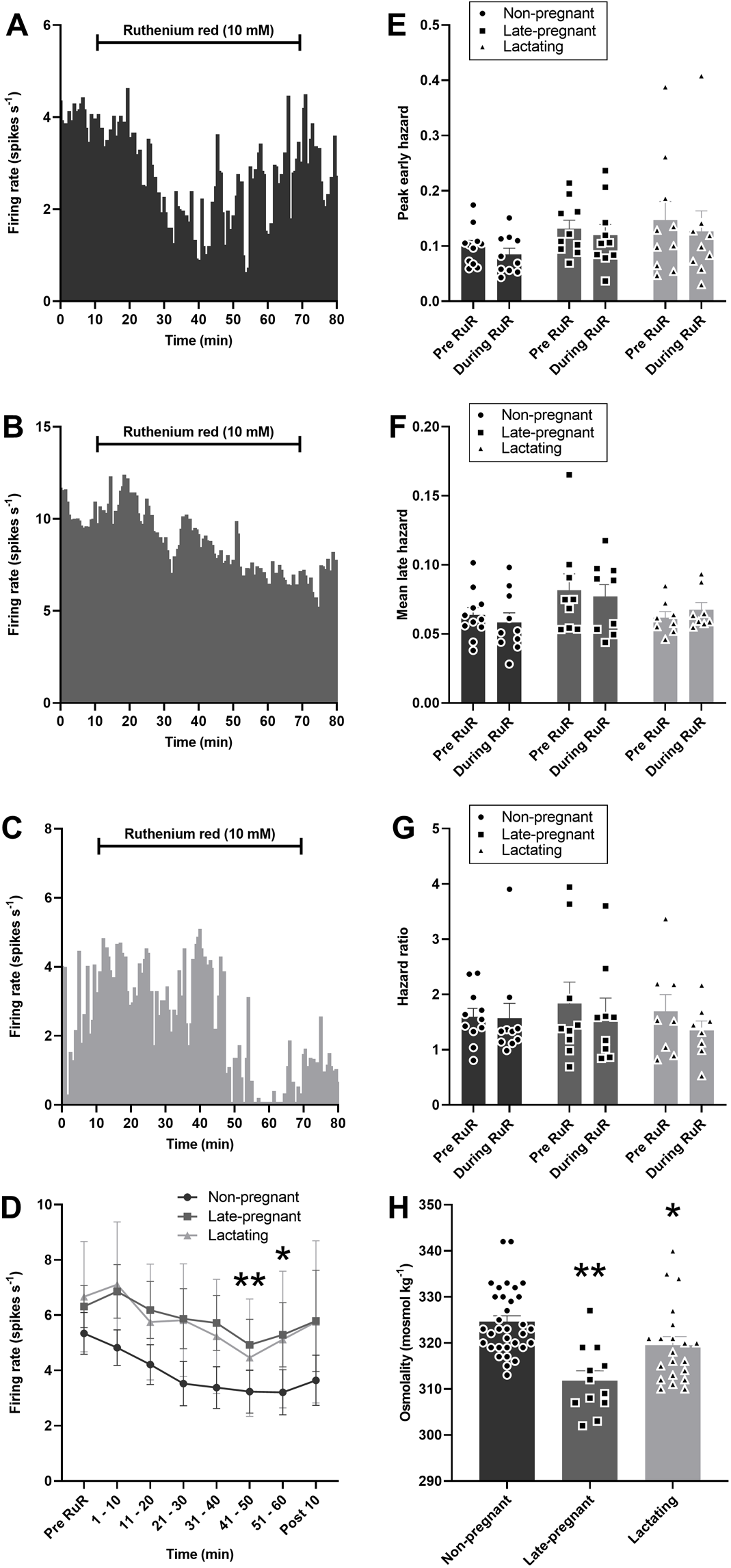
Ruthenium red inhibits vasopressin neuron activity in non-pregnant, late-pregnant and lactating rats. **A – C**. Ratemeter recordings (in 30 s bins) of the firing rates of vasopressin neurons in non-pregnant (A), late-pregnant (B) and lactating (C) rats before and during microdialysis administration of ruthenium red (RuR; 10 mM). **D**. Mean firing rate (in 10 min bins ± SEM) of vasopressin neurons in non-pregnant, late-pregnant and lactating rats before and during microdialysis administration of ruthenium red (10 mM). Ruthenium red induced a similar reduction in the firing rate of vasopressin neurons in non-pregnant, late-pregnant and lactating rats (REPRODUCTIVE STATUS: F_2,28_ = 0.69, P = 0.51; TIME: F_6,28_ = 5.94, P < 0.001; interaction between REPRODUCTIVE STATUS and TIME: F_12,32_ = 0.29, P = 0.99, two-way repeated measures ANOVA). *P < 0.05 and **P <0.01 compared to pre-RuR within TIME, Holm-Sidak post hoc tests. **E – G**. Mean (± SEM) peak early hazard (E), late hazard (F) and peak early:mean late hazard ratio (G) in vasopressin neurons from non-pregnant, late-pregnant and lactating rats. There was no effect of REPRODUCTUVE STATUS or TIME on the peak early hazard (F_1,28_ = 1.70, P = 0.20), mean late hazard (F_1,26_ = 0.37, P = 0.85) or hazard ratio (F_1,26_ = 0.21, P = 0.81) of vasopressin neurons during ruthenium red administration and no interaction between REPRODUCTUVE STATUS and TIME. **H**. Mean (± SEM) plasma osmolality in non-pregnant, late-pregnant and lactating rats. Plasma osmolality was affected by reproductive status (H = 20.0, P < 0.001). *P < 0.05 and ***P < 0.001, Dunn’s post hoc tests compared to non-pregnant rats.

By contrast to vasopressin neurons, oxytocin neuron activity was not affected by ruthenium red in non-pregnant or late-pregnant rats (F_1,9_ = 0.03, P = 0.87 for the main effect of REPRODUCTIVE STATUS; F_6,9_ = 1.10, P = 0.37 for the main effect of TIME; F_6,9_ = 1.00, P = 0.41 for the interaction, two-way repeated measures ANOVA; Figure 7). There was no effect of ruthenium red on the peak early hazard (F_1,9_ = 0.58, P = 0.46), mean late hazard (F_1,9_ = 1.97, P = 0.19) or hazard ratio (F_1,9_ = 0.09, P = 0.77) of oxytocin neurons and no interaction of ruthenium red with reproductive status (Figure 7).

**Figure 7.**
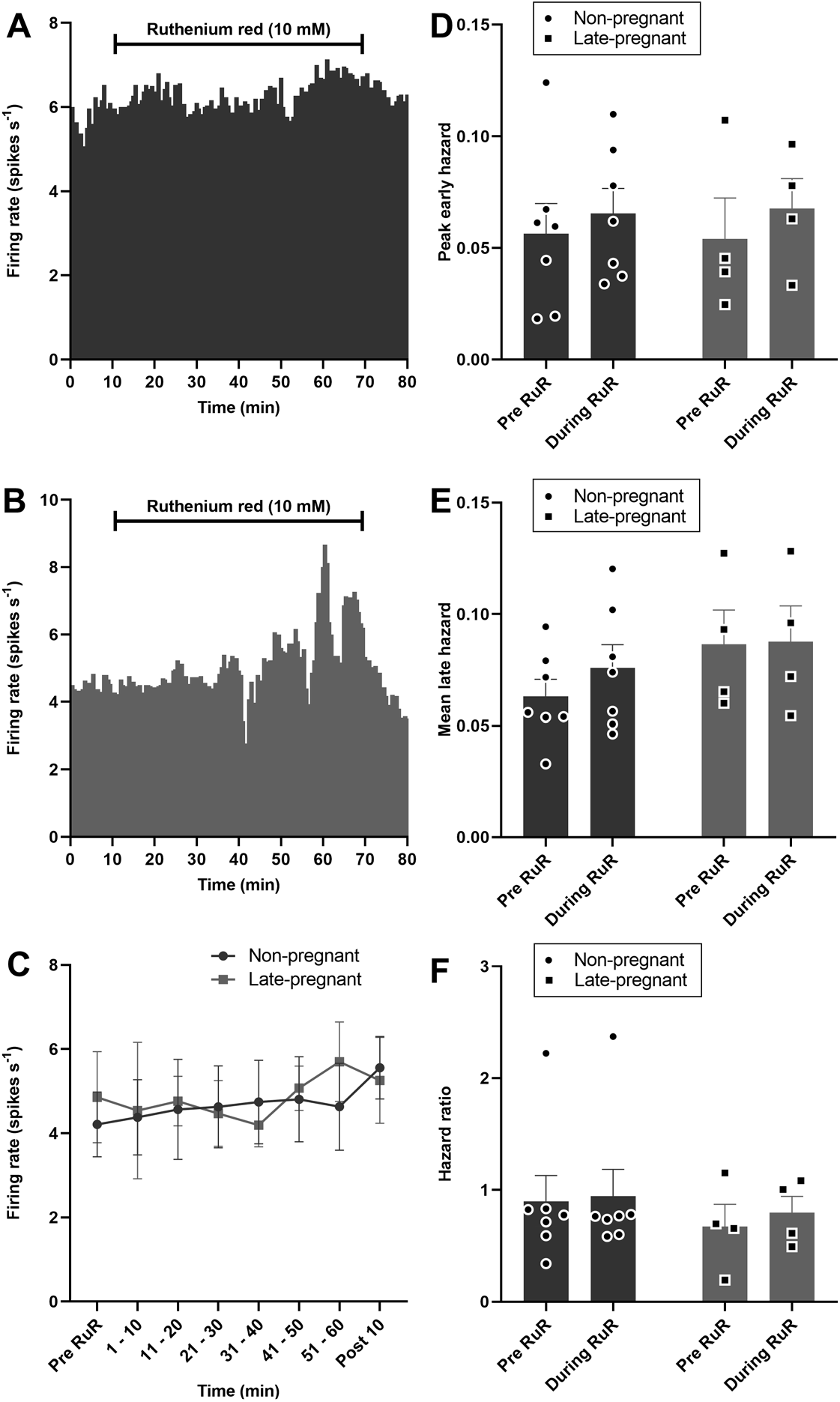
Ruthenium red does not affect oxytocin neuron activity in non-pregnant and late-pregnant rats. **A and B**. Ratemeter recordings (in 30 s bins) of the firing rates of oxytocin neurons in non-pregnant (A) and late-pregnant (B) rats before and during microdialysis administration of ruthenium red (RuR; 10 mM). **C**. Mean firing rate (in 10 min bins ± SEM) of oxytocin neurons in non-pregnant, and late-pregnant rats before and during microdialysis administration of ruthenium red (10 mM). Oxytocin neuron activity was not affected by ruthenium red in non-pregnant or late-pregnant rats (REPRODUCTIVE STATUS: F_1,9_ = 0.03, P = 0.87; TIME: F_6,9_ = 1.10, P = 0.37; interaction between REPRODUCTIVE STATUS and TIME: F_6,9_ = 1.00, P = 0.41, two-way repeated measures ANOVA). **D – F**. Mean (± SEM) peak early hazard (D), late hazard (E) and peak early:mean late hazard ratio (F) in oxytocin neurons from non-pregnant, late-pregnant and lactating rats. There was no effect of REPRODUCTIVE STATUS or TIME on the peak early hazard (F_1,9_ = 0.58, P = 0.46), mean late hazard (F_1,9_ = 1.97, P = 0.19) or hazard ratio (F_1,9_ = 0.09, P = 0.77) of oxytocin neurons during ruthenium red administration and no interaction between REPRODUCTUVE STATUS and TIME.

### ΔN-TRPV1 mRNA and vasopressin mRNA expression in the supraoptic nucleus of non-pregnant and late-pregnant rats

Labelling of ΔN-TRPV1 mRNA and vasopressin mRNA was most prominent within the supraoptic nucleus (Supplementary Figure 1). ΔN-TRPV1 mRNA was evident as punctate labelling whereas vasopressin mRNA labelling was evident throughout the cytoplasm in the cell bodies. There was no difference in ΔN-TRPV1 mRNA expression (P = 0.66, unpaired t-test) or vasopressin mRNA expression (P = 0.71) between non-pregnant and late-pregnant rats. Co-expression of ΔN-TRPV1 mRNA with vasopressin mRNA was not different between non-pregnant and late-pregnant rats (P = 0.41; Figure 8). In addition, co-expression of vasopressin mRNA with ΔN-TRPV1 mRNA was not different between non-pregnant (50.9 ± 2.5%) and late-pregnant rats (46.2 ± 0.3%; P = 0.63, unpaired t-test).

**Figure 8.**
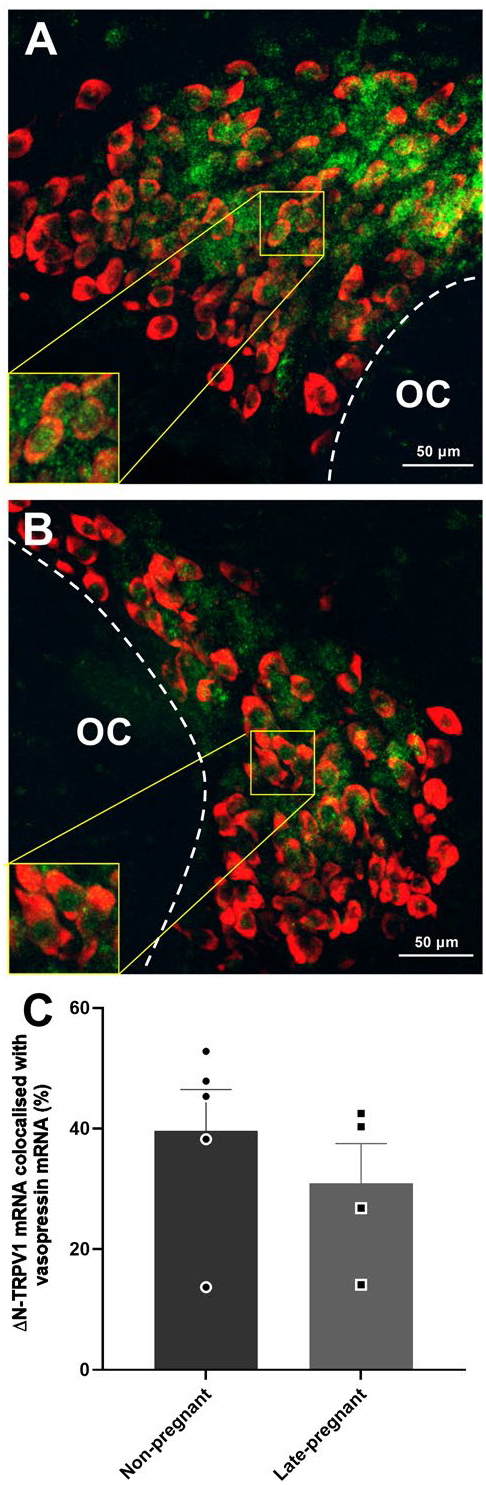
Vasopressin mRNA and ΔN-TRPV1 mRNA expression in the supraoptic nucleus of non-pregnant and late-pregnant. **A and B**. Example compressed Z-stacks of vasopressin mRNA (red) and ΔN-TRPV1 mRNA (green) labelling in the supraoptic nucleus of non-pregnant (A) and late-pregnant (B) rats. OC: optic chiasm. **C**. Mean levels of ΔN-TRPV1 mRNA and vasopressin mRNA co-labelling in the supraoptic nucleus of non-pregnant and late-pregnant rats. There was no difference in ΔN-TRPV1mRNA expression (P = 0.66, unpaired t-test), vasopressin mRNA expression (P = 0.71), or co-expression of ΔN-TRPV1 mRNA and vasopressin mRNA between non-pregnant and late-pregnant rats (P = 0.41).

## Discussion

Here, we found that basal firing rate was not different between oxytocin neurons in non-pregnant, late-pregnant and lactating rats or between vasopressin neurons in non-pregnant, late-pregnant and lactating rats, consistent with our previous observations (24, 28). Nevertheless, hazard analyses revealed a reorganisation of activity in oxytocin neurons, but not vasopressin neurons, in late-pregnant and lactating rats that favoured the occurrence of short intervals between successive spikes that are necessary to achieve the high firing rates evident during burst firing in birth and lactation. By contrast to oxytocin neurons, the activity of vasopressin neurons in late-pregnant and lactating rats was indistinguishable from that in non-pregnant rats. The maintenance of vasopressin neuron activity in the face of lower plasma osmolality in late pregnancy appears to involve increased intrinsic osmosensitivity.

### Increased post-spike hyperexcitability in oxytocin neurons in late pregnancy and lactation

Hazard functions calculate the probability of the neuron firing a subsequent spike in any given interval following a preceding spike, which reflects post-spike excitability. Changes in the hazard function are inferred to result from changes in membrane potential after each spike and therefore reveal the influence of post-spike potentials on activity patterning (29). The measures derived from the hazard function to describe post-spike excitability are the peak early hazard, the mean late hazard and the peak early:mean late hazard ratio. The peak early hazard is inferred to reflect the transient impact of the mAHP and sADP superimposed on baseline excitability (29) but can also be influenced by other transient changes in membrane potential, such as that induced by the transient outward rectifier (30). The mean late hazard reflects the steady-state excitability of neurons and is inferred to reflect the influence of ongoing synaptic input and underlying membrane potential (29). The peak early:mean late hazard ratio quantifies the shape of the hazard function by scaling the peak early hazard to the mean late hazard to reveal the relative influence of post-spike potentials and ongoing synaptic input/baseline membrane potential to spike firing.

The oxytocin system undergoes extensive plasticity during pregnancy to prepare for birth and lactation (31) that includes the emergence of an sADP in oxytocin neurons, as well as a shorter mAHP (32, 33). The sADP supports burst firing by increasing post-spike excitability in vasopressin neurons (34, 35). Hence, the emergence of an sADP in late pregnancy would be expected to increase post-spike excitability of oxytocin neurons to support burst firing essential for normal birth and lactation. While we did not find an overall change in post-spike excitability across all oxytocin neurons, there was clear post-spike hyperexcitability in a similar proportion of oxytocin neurons to those that express a measurable sADP in late pregnancy and lactation (32, 33). Hence, the simplest explanation of increased incidence of post-spike hyperexcitability observed here is an enhanced expression of the sADP by some oxytocin neurons.

Oxytocin generally express a sADP under basal conditions but this is usually masked by the occurrence of a larger mAHP (16). Hence, the emergence of post-spike hyperexcitability in oxytocin neurons might reflect a reduced influence of the mAHP that unmasks the sADP. The mAHP increases the regularity of oxytocin neuron firing during basal activity (36) and so a reduced mAHP could contribute to the increased variability of spike firing (CV) observed here. Increased variability of spike firing is correlated with burst firing in oxytocin neurons (37, 38). The observed positive correlation of CV with hazard ratio suggests that increased post-spike hyperexcitability might contribute to increased variability of firing. If this is the case, then the oxytocin neurons with higher post-spike hyperexcitability might be those that are most likely to adopt bursting activity (39).

### Increased excitability in vasopressin neurons in late pregnancy and lactation

Vasopressin secretion is principally stimulated by body fluid osmolality, and so lowered osmolality normally decreases vasopressin secretion (3). However, despite water retention profoundly reducing osmolality in pregnancy and lactation (40), vasopressin levels are unchanged, or even increased in pregnant and lactating humans and rats (4, 5, 41–44).

Under basal conditions, osmolality directly excites vasopressin neurons (11) via ΔN-TRPV1 in their plasma membrane (45, 46). ΔN-TRPV1 inhibition with ruthenium red reduced vasopressin neuron activity in non-pregnant rats, showing for the first time that activation of ΔN-TRPV1 contributes to the basal activity of vasopressin neurons *in vivo*. Surprisingly, the activity of oxytocin neurons was not affected by ruthenium despite oxytocin neurons being as sensitive to excitation by acute osmotic stimulation as vasopressin neurons (47). Hence, osmosensitivity of oxytocin neurons appears to be less dependent on ΔN-TRPV1 than for vasopressin neurons.

While ruthenium red inhibits ΔN-TRPV1 (48), it is a broad-spectrum TRPV inhibitor and has several other well-characterised effects, including reduction of calcium release from the endoplasmic reticulum via ryanodine receptor inhibition (49). However, ruthenium red is relatively membrane impermeable and is therefore administered directly into cells to study its effects on intracellular calcium release, for example, by inclusion in the micropipette solution during whole-cell patch clamp recording (50). Intracellular administration of 20 µM ruthenium red is required to inhibit the ryanodine receptor in supraoptic nucleus neurones (51), whereas an extracellular concentration of 10 µM is required to inhibit ΔN-TRPV1 channels on supraoptic nucleus neurones *in vitro* (45). Here, ruthenium red was administered at 10 mM in the dialysate and microdialysis administration is estimated to deliver a 1000-fold lower drug concentration to the supraoptic nucleus than is contained in the dialysate (52). Hence, the extracellular concentration of ruthenium red in the supraoptic nucleus in our experiments should be ∼10 µM, which should be sufficient to inhibit ΔN-TRPV1 channels on the cell surface but not ryanodine receptors on the endoplasmic reticulum. Indeed, inhibition of intracellular calcium release by ruthenium red reduces sADP amplitude in supraoptic nucleus neurons (51) but ruthenium red did not affect the shape of hazard function in the current experiments, confirming that it did not inhibit calcium release from intracellular stores. Rather, ruthenium red caused a generalised reduction in post-spike excitability that would be expected to result from sustained hyperpolarisation of membrane potential (29).

Ruthenium red reduced vasopressin neuron activity to the same extent in late-pregnant and lactating rats as in non-pregnant rats, indicating that the ΔN-TRPV1 contribution to basal vasopressin neuron activity is maintained at lower osmolality in late-pregnancy and lactation. ΔN-TRPV1 mRNA expression was not different between the supraoptic nuclei of non-pregnant and late-pregnant rats, suggesting that ΔN-TRPV1 synthesis was similar. Hence, increased ΔN-TRPV1 function might underpin the lower osmotic threshold for vasopressin secretion that develops in pregnancy (53) and thereby contribute to the expansion of blood volume required to cope with the cardiovascular demands of pregnancy and lactation. TRPV2 and TRPV4 are expressed by vasopressin neurons (54) and each can dimerise with TRPV1 (55, 56). We are currently investigating whether TRPV4 dimerisation with ΔN-TRPV1 might be able to underpin increased osmosensitivity of vasopressin neurons in late pregnancy and lactation.

Approximately half of the ΔN-TRPV1 mRNA was expressed in vasopressin neurons in non-pregnant and late-pregnant rats. Therefore, the remaining ΔN-TRPV1 mRNA must be expressed by other cells in the supraoptic nucleus, presumably oxytocin neurons. Hence, the expression of ΔN-TRPV1 mRNA is unlikely to underpin the lower sensitivity of oxytocin neurons to ruthenium red inhibition than vasopressin neurons.

## Conclusion

Taken together, the current data suggest that increased intrinsic excitability of oxytocin and vasopressin neurons contribute to normal pregnancy and lactation. Increased post-spike hyperexcitability of oxytocin neurons might help underpin burst firing required for delivery of the fetus during birth and milk ejection during suckling, while increased intrinsic osmosensitivity of vasopressin neurons resets the osmotic threshold for vasopressin secretion to drive blood volume expansion during pregnancy.

## Acknowledgements

This work was supported by project grants from the Marsden Fund of the Royal Society of New Zealand and the New Zealand Health Research Council (CHB), a University of Otago Research Grant (CHB) and by University of Otago Doctoral Scholarships (MRP and EFB). We thank Ms Mehwish Abbasi, Ms Zoë Jacquiery and Dr Victoria Scott for recordings of basal activity in non-pregnant, late-pregnant and lactating rats.

## Author contributions

RAA and CHB designed the electrophysiology study. MRP, AJS and CHB completed the electrophysiology experiments and analyses. GTB and CHB designed the RNAscope study. GTB, EFB and IC completed and analysed the RNAscope experiment. All authors contributed to the interpretation of the results. MRP and CHB wrote the first draft manuscript and all authors provided intellectual input for the final version.

---

**Supplementary Figure 1.**
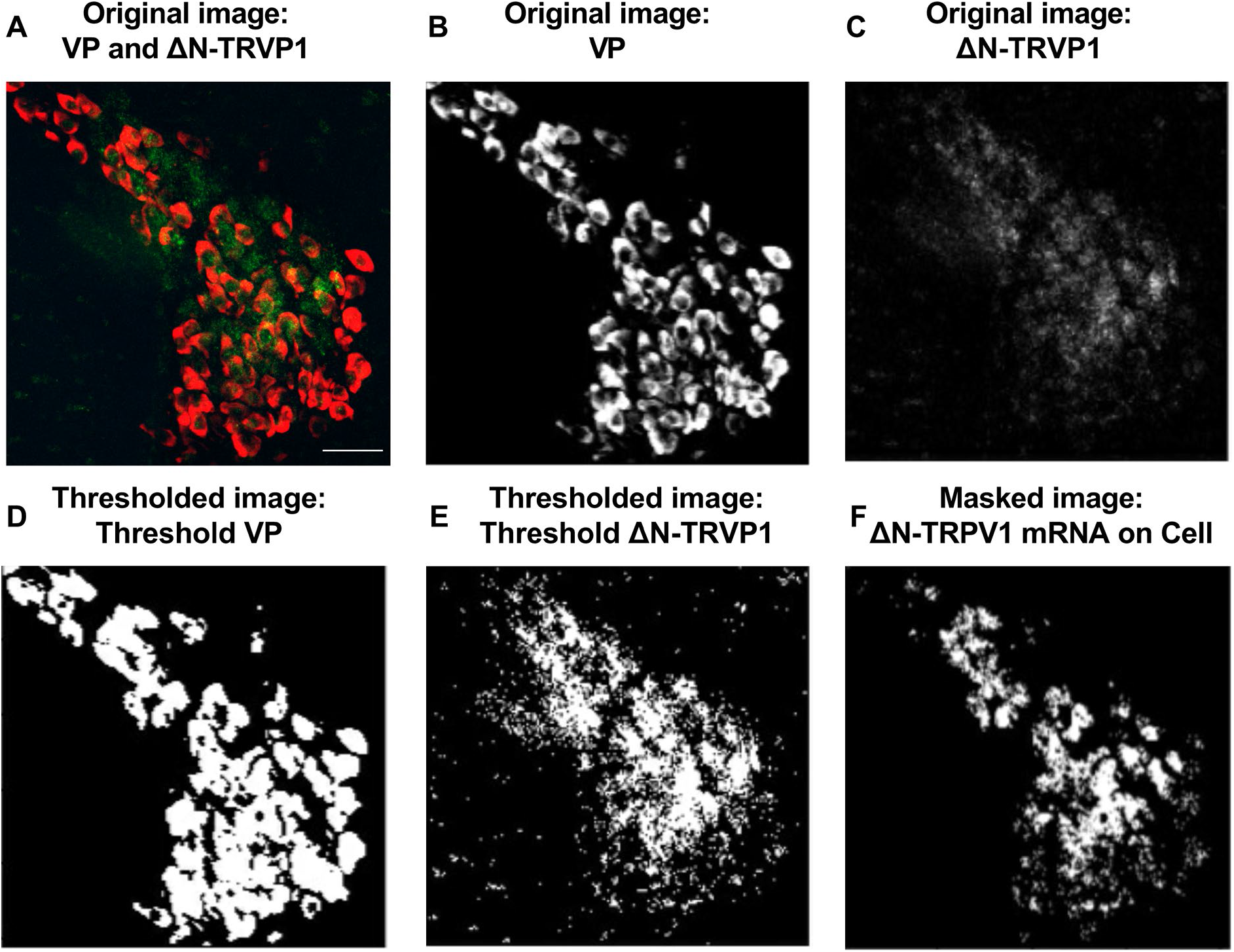
Example of the analysis of vasopressin mRNA and ΔN-TRPV1 mRNA expression in the supraoptic nucleus of non-pregnant and late-pregnant rats. **A**. Example micrograph of vasopressin mRNA and ΔN-TRPV1 mRNA labelling in the supraoptic nucleus as shown in Figure 8B, generated by compression of Z-stacks using ImageJ Fiji Z-projection Max intensity. **B and C**. Images generated by splitting the image in panel A into the vasopressin mRNA channel (B) and ΔN-TRPV1 mRNA channel (C). **D and E**. Thresholded images for vasopressin mRNA expression from panel B and ΔN-TRPV1 mRNA from panel C, generated using Cell Profiler™. **F**. Masked image created by overlaying the two thresholded images in panels D and E, showing only pixels that were co-labelled for vasopressin mRNA and ΔN-TRPV1 mRNA. Note that vasopressin mRNA labelling did not allow differentiation between the borders of individual cells in all sections. Therefore, it was not possible to quantify of numbers of ΔN-TRPV1 mRNA-positive vasopressin neurons.

**Supplementary Figure 2.**
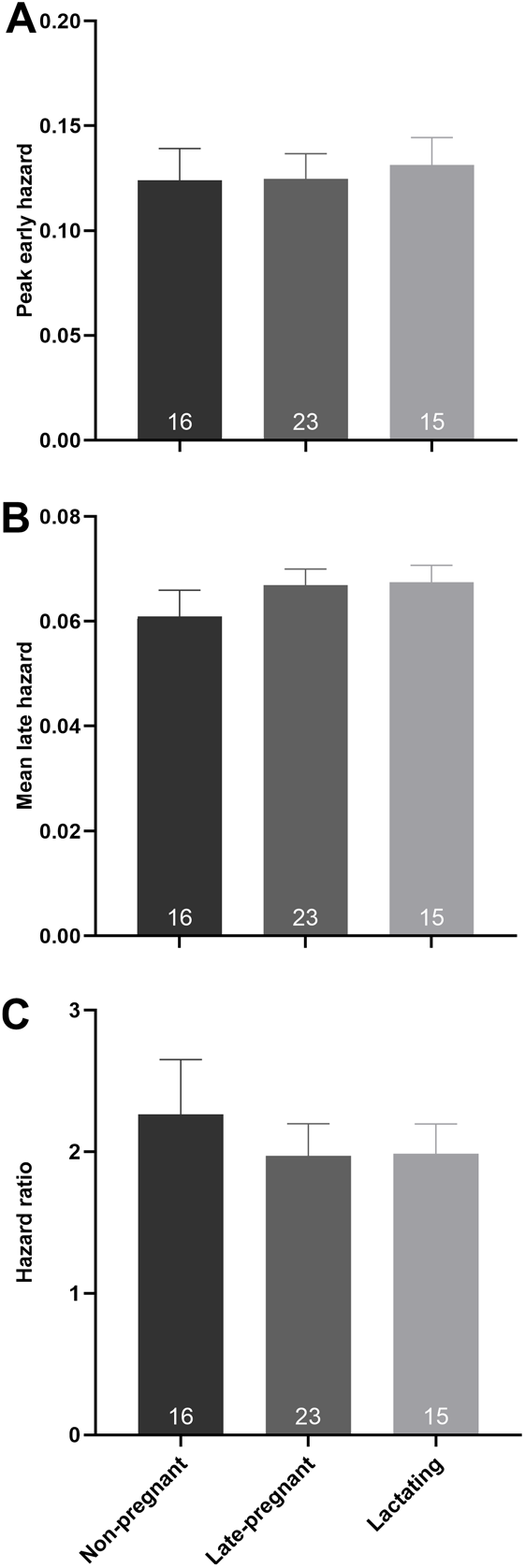
**A – C**. Mean (± SEM) peak early hazard (A), mean late hazard (B) and peak early:mean late hazard ratio (C) of phasic vasopressin neurons in non-pregnant, late-pregnant and lactating rats. There was no effect of reproductive status on the peak early hazard (H = 0.17, P = 0.92, one-way ANOVA on ranks), mean late hazard (H = 4.27, P = 0.12) or hazard ratio (H = 0.41, P = 0.82) of phasic vasopressin neurons in non-pregnant, late-pregnant and lactating rats.

**Supplementary Figure 3.**
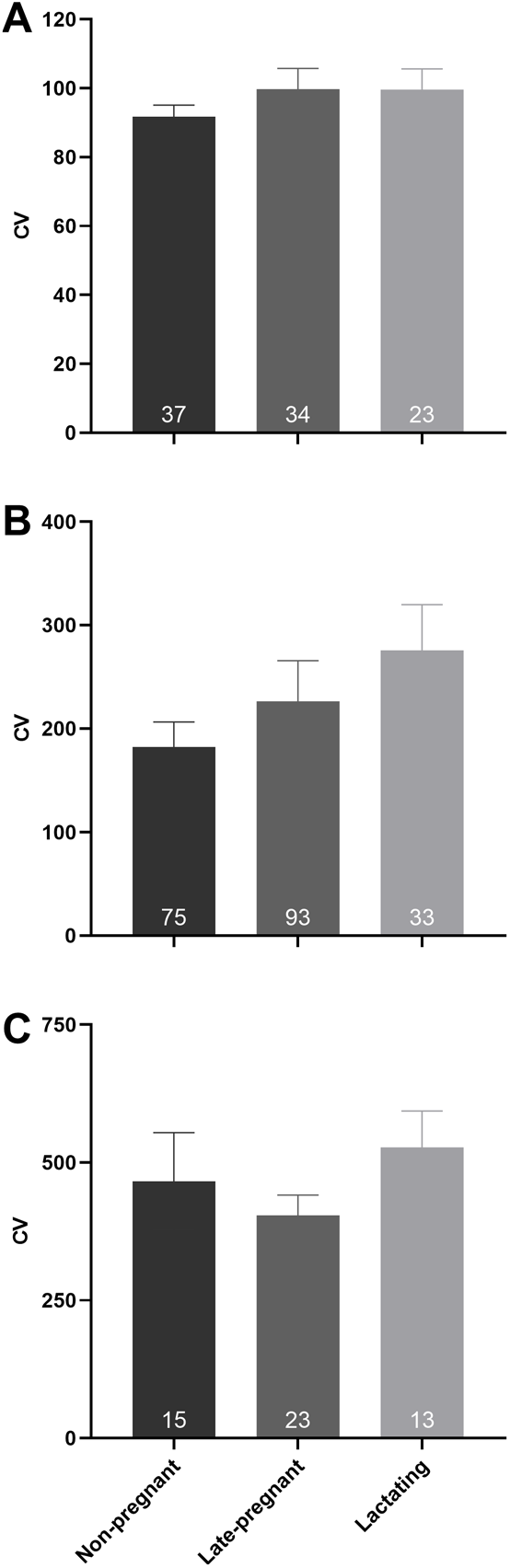
**A – C**. Mean (± SEM) coefficient of variation of inter-spike interval of oxytocin neurons (A), vasopressin neurons (B) and phasic vasopressin (C) neurons. There was no effect of reproductive status on the CV of oxytocin neurons (H = 0.76, P = 0.69, one-way ANOVA on ranks), vasopressin neurons (H = 5.43, P = 0.07), or phasic vasopressin neurons (H = 2.73, P = 0.25).

